# Vimentin filaments interact with the mitotic cortex allowing normal cell division

**DOI:** 10.1101/356642

**Authors:** Sofia Duarte, Álvaro Viedma-Poyatos, Elena Navarro-Carrasco, Alma E. Martínez, María A. Pajares, Dolores Pérez-Sala

**Author notes:** To whom correspondence should be addressed at: Department of Structural and Chemical Biology Centro de Investigaciones Biológicas Consejo Superior de Investigaciones Científicas (C.S.I.C.) Ramiro de Maeztu, 9 28040 Madrid, Spain. Present address: Centro Cardiologico Monzino, Via Carlo Parea, 4, 20138 Milano MI, Italia.

## Abstract

The vimentin network displays remarkable plasticity to support basic cellular functions. Here, we show that in several cell types vimentin filaments redistribute to the cell periphery during mitosis, forming a robust scaffold interwoven with cortical actin and affecting the mitotic cortex properties. Importantly, the intrinsically disordered tail domain of vimentin is essential for this redistribution, which allows normal mitotic progression. A tailless vimentin mutant forms curly bundles, which remain entangled with dividing chromosomes leading to mitotic catastrophes or asymmetric partitions. Serial deletions of the tail domain induce increasing impairments of cortical association and mitosis progression. Disruption of actin, but not of microtubules, mimics the impact of tail deletion. Pathophysiological stimuli, including HIV-protease and lipoxidation, induce similar alterations. Interestingly, filament integrity is dispensable for cortical association, which also occurs in vimentin particles. These results unveil novel implications of vimentin dynamics in cell division by means of its interplay with the mitotic cortex.

## Introduction

The vimentin filament network provides architectural support for cells and contributes to the positioning and function of cellular organelles ^1-3^. Vimentin plays multiple roles in cell regulation by interacting with signaling proteins, adhesion molecules ^4,5^, chaperones ^6,7^ and other cytoskeletal elements ^8,9^. The vimentin monomer consists of 466 residues organized in a central rod of predominantly α-helical structure flanked by intrinsically disordered N- and C-terminal domains (Fig. 1A). Vimentin polymerization is believed to progress from parallel dimers to antiparallel tetramers, eight of which associate laterally in “unit length filaments” that engage head to tail to form filaments. The vimentin network is highly dynamic and rapidly responds to heat-shock, oxidative and electrophilic stresses, ATP and divalent cation availability ^10-12^, playing a key role in cell adaptation.

**Fig. 1.**
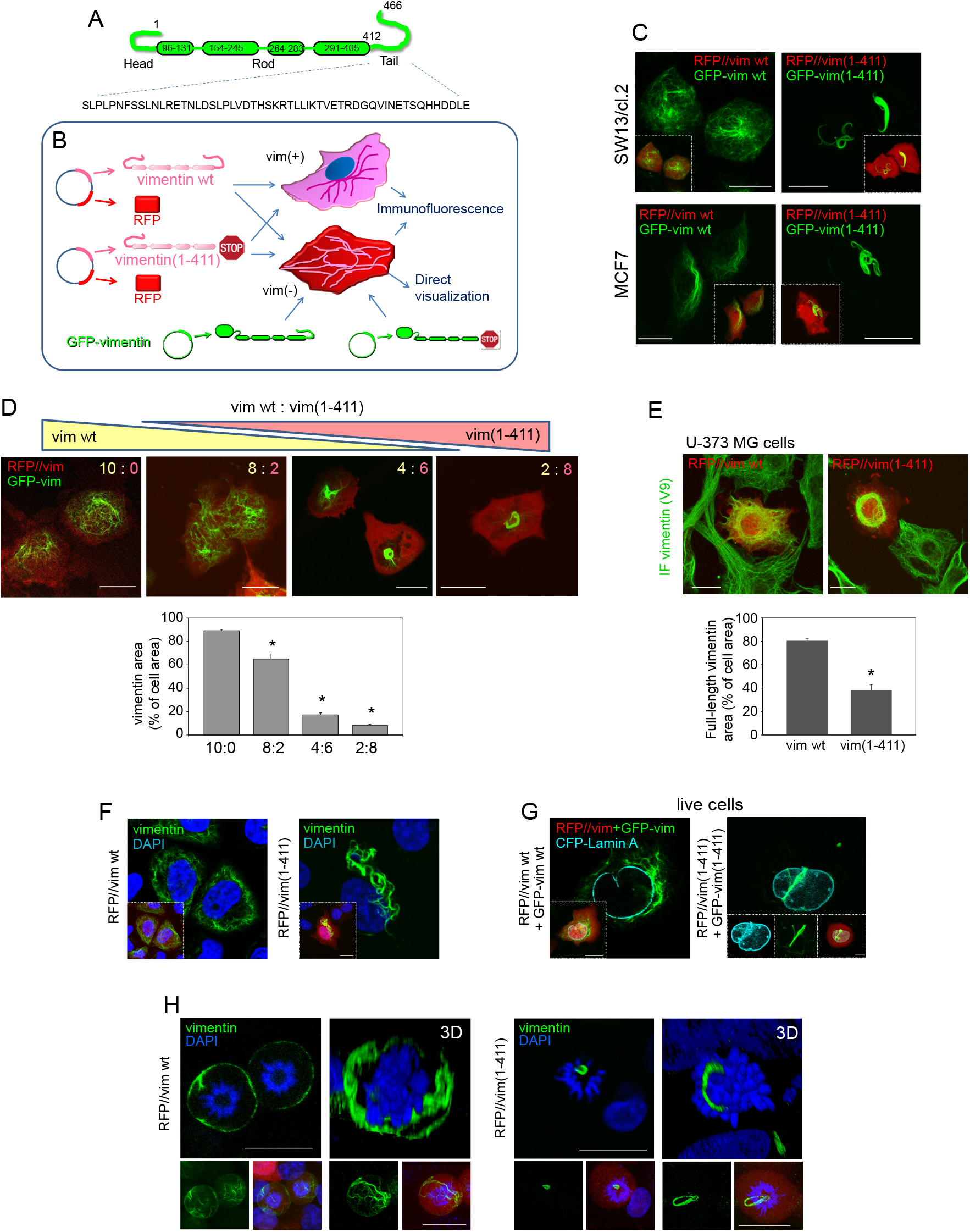
Tailless vimentin(1-411) disrupts wt vimentin distribution and interferes with chromosomes in mitosis. (A) Schematic view of vimentin domains. Residues are numbered. The tail domain sequence is displayed in full. (B) Scheme showing the experimental strategies: bicistronic plasmids coding for DsRed2 fluorescent protein (RFP) and vimentin wt (RFP//vim wt) or tailless (residues 1-411) (RFP//vim(1-411)) were transfected into vimentin expressing cells (vim +) or vimentin-deficient cells (vim -), alone, for detection by immunofluorescence, or together with a small amount of the corresponding GFP-vimentin construct (GFP-vim) for direct visualization. (C) SW13/cl.2 human adrenocarcinoma or MCF7 breast carcinoma cells transfected with the indicated constructs were observed live 48 h later. (D) SW13/cl.2 cells were transfected with different proportions of constructs coding for vimentin wild type (wt, yellow) or tailless (1-411, pink), as detailed in Materials and Methods. Vimentin condensation was measured as the area occupied by vimentin (green fluorescence) with respect to the total cellular area (red background). The histogram shows average values ± SEM of twenty determinations. \**P*<0.05 by Student’s *t*-test. (E) U-373 MG astrocytoma cells were transfected with RFP//vimentin wt or RFP//vimentin(1-411). Full-length vimentin condensation was assessed by immunofluorescence with V9 anti-vimentin antibody, which recognizes the tail domain (green), and estimated as the ratio between the green signal area and the total cell area (red). The histogram shows average values ± SEM of twenty determinations. \**P*<0.001 by Student’s *t*-test. (F) SW13/cl.2 cells were transfected with RFP//vimentin wt or (1-411) and vimentin distribution was assessed by immunofluorescence. Nuclei were counterstained with DAPI and single overlay sections are shown. (G) Cells were visualized live after transfection with GFP-vimentin wt or GFP-vimentin(1-411) plus CFP-lamin A to delimit the nuclear envelope. Insets in (F) and (G) display overall projections of merged images or individual channels. (H) Cells were transfected with RFP//vimentin wt or (1-411) and vimentin distribution in mitosis was observed by immunofluorescence. Single sections taken at mid-cell height (left images) and 3D-projections (right images) are shown. Small panels below each image depict overall projections for vimentin alone (left) or for the three channels (vimentin, RFP and DAPI). Scale bars, 20 µm.

Posttranslational modifications are critical for fast and versatile network remodeling ^13^. Phosphorylation of specific residues regulates vimentin assembly and involvement in migration and invasion ^14,15^. In mitosis, vimentin phosphorylation is regulated in a spatio-temporal manner, leading to filament disassembly in certain cell types ^16,17^. Besides, oxidative and electrophilic modifications of vimentin’s single cysteine drastically alter network organization, highlighting its crucial role for filament architecture ^12,18^.

Vimentin is also the substrate for several proteases, the resulting fragments performing additional cellular roles. Calpains cleave vimentin N-terminus impairing polymerization ^19^, and calpain-truncated soluble vimentin associates with phosphorylated-ERK playing a role in axonal regeneration ^20^. Conversely, caspase-generated amino-terminal fragments exert pro-apoptotic effects ^21^. Vimentin C-terminus also contains sites for viral protease cleavage. Moloney mouse sarcoma virus infection produces fragments lacking all or part of the C-terminal tailt^22^, whereas the HIV-type 1 protease reportedly cleaves vimentin after L423 ^23^. Curiously, chemicals like gambogic acid promote vimentin cleavage in cells by yet unidentified proteases, rendering products missing sequences before S51 and/or after R424 ^24^.

The role of the tail domain in vimentin organization is still incompletely understood. Purified tailless vimentin (vimentin(1-411)) polymerizes in vitro into normal filaments ^25-27^, presenting oligomerization and sedimentation behaviors highly similar to those of full-length vimentin ^28^, although higher heterogeneity and wider average diameter ^29^ have also been noted. In turn, the vimentin tail has been suggested to undergo conformational changes during filament elongation and assembly in vitro ^30^, and to modulate interactions with divalent cations ^25,31^.

In cells, vimentin(1-411) mutants form either normal extended arrays or filaments with a tendency to collapse, depending on the experimental system ^26,27^. Additionally, the tail domain has been proposed to act as a cytoplasmic retention signal ^32^ and contribute to filament stability ^26^. However, the mechanism(s) by which C-terminally truncated vimentin forms induce cellular perturbations has not been fully elucidated.

Vimentin is an exquisite sensor for oxidative and electrophilic stresses ^12^, and presents sites for modification by electrophilic lipids (lipoxidation) across the whole monomer ^33^. While exploring these modifications, we have observed that C-terminal truncated mutants, particularly vimentin(1-411) and the reported product of HIV-protease cleavage ^23^, vimentin(1-423), exert deleterious cellular effects, with formation of juxtanuclear bundles and aberrant mitosis. A deeper analysis showed that, if not disassembled, full-length vimentin redistributed to the cell periphery in mitosis, in close interplay with the mitotic cortex, leaving ample space for dividing chromosomes. Strikingly, vimentin(1-411) did not reach the cell periphery and snared the mitotic apparatus. These observations identify vimentin as a novel element of the mitotic cortex in a tail domain-dependent manner.

## Results

### 1. Vimentin tail is necessary for appropriate network distribution in resting and mitotic cells

To elucidate the roles of the vimentin tail domain in filament assembly and stress responses, we employed several strategies to express wild type (wt) or vimentin(1-411) in vimentin-positive or -deficient cells (Fig. 1B). Vimentin-deficient adrenocarcinoma SW13/cl.2 or breast carcinoma MCF7 cells were co-transfected with RFP//vimentin bicistronic plasmids plus GFP-vimentin vectors for network visualization in live cells. Whereas vimentin wt formed an extended network, the organization of vimentin(1-411) was drastically altered, forming curly juxtanuclear filament bundles (Fig. 1C). Importantly, transfecting vimentin(1-411) in excess over wt, impaired network extension and induced its condensation in coiled bundles (Fig. 1D). Moreover, in cells expressing endogenous vimentin, overexpression of vimentin(1-411) but not vimentin wt, markedly disrupted network distribution, causing filament retraction from the cell periphery and perinuclear condensation (quantitated in Fig. 1E). Therefore, although vimentin(1-411) polymerization is not impeded, its cellular organization is severely altered. Moreover, vimentin(1-411) exerts deleterious effects on the organization of full-length vimentin, which depend on the proportion of both forms.

Detailed observation of nuclear structures showed that while vimentin wt filaments extended outwards from the nuclear periphery, vimentin(1-411) thick bundles displayed extensions into the area covered by DAPI staining (Fig. 1F), appearing in deep invaginations of the nuclear envelope or distributing between nuclear lobules (Fig. 1G and Supplementary Fig. 1). This drove us to assess vimentin(1-411) behavior in mitotic cells. In SW13/cl.2 cells vimentin wt did not disassemble in mitosis but remained as robust filaments with a marked peripheral distribution (Fig. 1H). In sharp contrast, vimentin(1-411) remained tightly packed in coiled bundles, frequently in close proximity of condensed chromosomes or forming loops encircling them (Fig. 1H and Supplementary Fig. 1).

### 2. Expression of tailless vimentin leads to aberrant mitosis

The striking pattern of vimentin(1-411) prompted us to monitor mitotic progression. Time-lapse experiments showed that cells expressing vimentin wt divided regularly, with an interval between cell rounding and daughter cell separation of 1-2 h (Fig. 2A and Supplementary videos 1 and 2), generally yielding homogeneous vimentin distribution. Conversely, cells harboring vimentin(1-411) bundles suffered diverse perturbations. Some cells attempted to divide for several hours and died during or shortly after mitosis, undergoing extensive membrane blebbing, typical of mitotic catastrophe (Fig. 2A, middle panels, and Supplementary video 3). Alternatively, some cells successfully completed division through the asymmetric partitioning of vimentin, implying retention of vimentin(1-411) in one daughter cell and “rejuvenation” of the other by initial elimination of vimentin coils (Fig. 2A, lower panels, and Supplementary video 4). Thus, the tail domain is necessary for normal vimentin dynamics during mitosis (schematized in Fig. 2B).

**Fig. 2.**
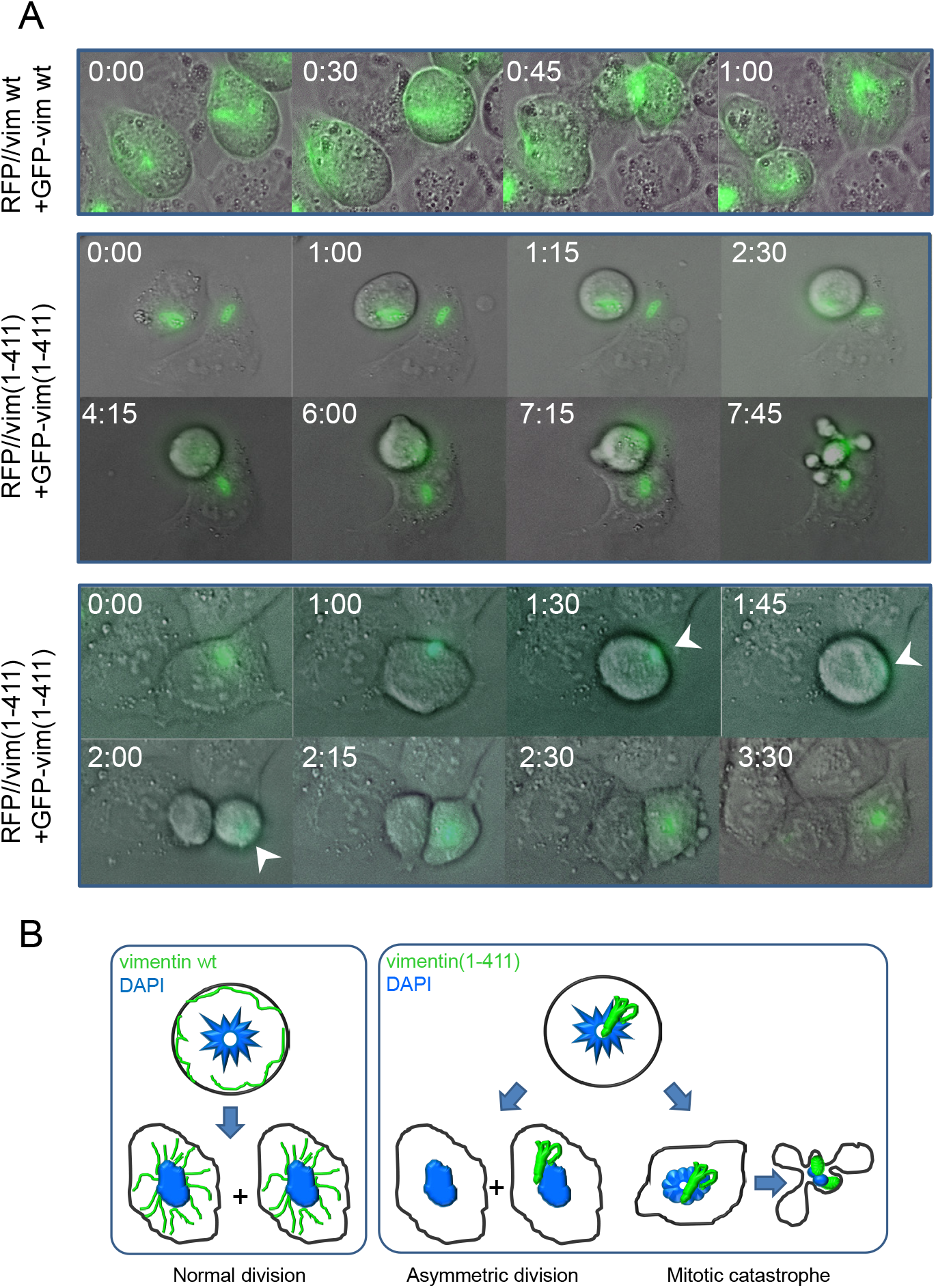
Live cell monitoring of cells expressing vimentin wt or vimentin(1-411) during mitosis. (A) SW13/cl.2 cells were transfected with RFP//vimentin plus GFP-vimentin wt or (1-411), or the equivalent constructs for vimentin(1-411), as indicated, for live cell monitoring by time-lapse microscopy. Several fields were randomly selected and images were acquired every 15 min. Representative images of the overlays of DIC and green fluorescence at the indicated time points are shown. (B) Schematic representation of the main fates observed for cells transfected with each construct.

### 3. Full-length vimentin but not vimentin(1-411) localizes to the mitotic cortex in an actin-dependent manner

Vimentin filaments maintain a close crosstalk with microtubules and microfilaments, which influence vimentin distribution. The interplay of vimentin wt or (1-411) with tubulin and actin was analyzed by disruption of these structures with various agents (Fig. 3A). Vimentin wt partially co-localized with tubulin in interphase, whereas in mitosis, vimentin underwent peripheral redistribution and tubulin concentrated at the mitotic spindle (Fig. 3B). Vimentin(1-411) bundles were largely unconnected to microtubules in resting cells, but remained adjacent to the mitotic spindle during cell division, frequently interfering with the position of chromosomes or microtubules (Fig. 3B). Acute microtubule disruption with nocodazole completely blocked the formation of the mitotic spindle and caused misorientation of chromosomes, without altering the peripheral distribution of vimentin wt in mitosis or the presence of vimentin(1-411) bundles (Fig. 3B).

**Fig. 3.**
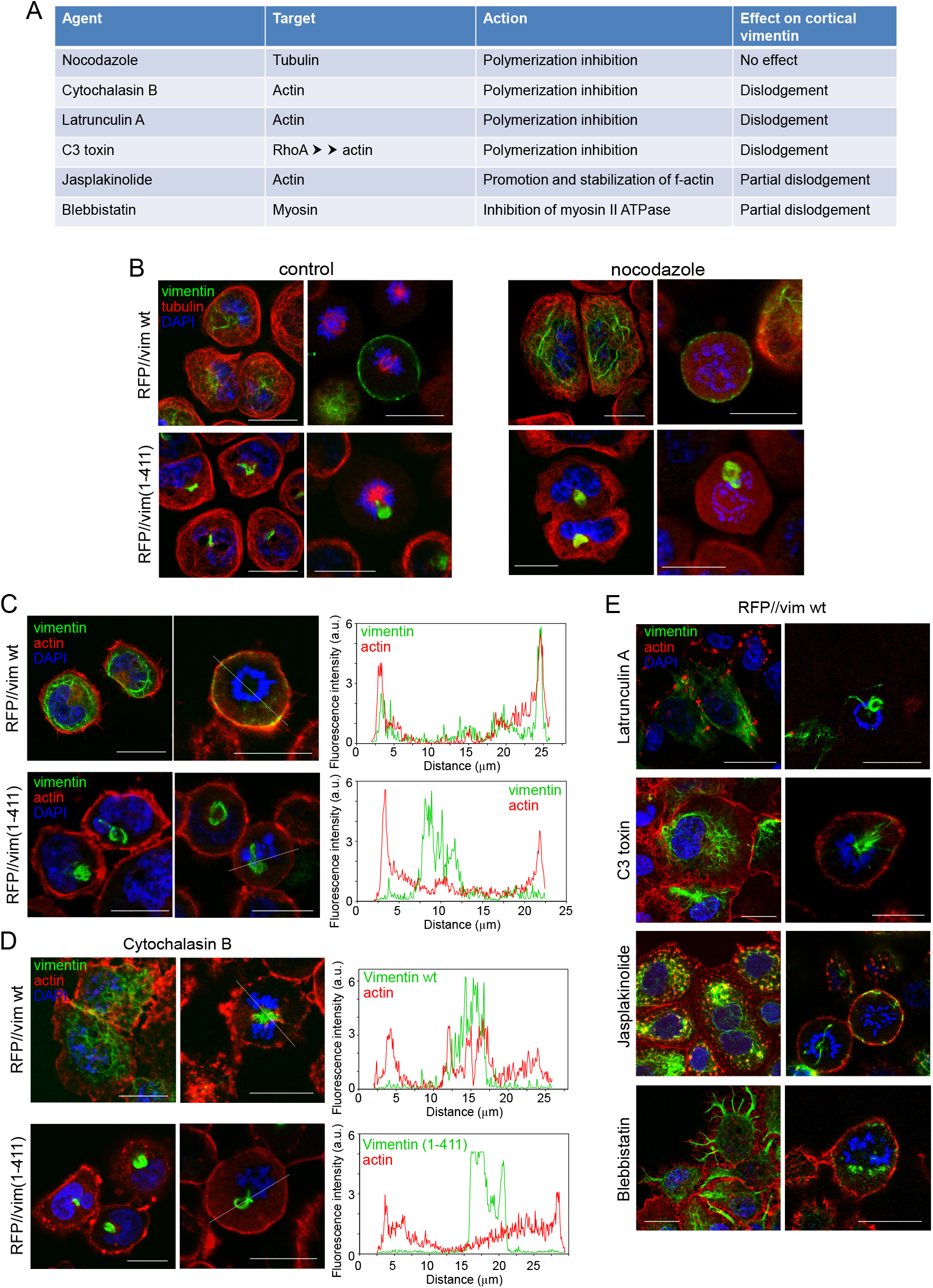
Effect of disrupting microtubules or actin filaments on the distribution of vimentin wt and vimentin(1-411) (A) Agents used to disrupt cytoskeletal structures. (B-E) The distribution of vimentin and tubulin or actin was assessed in interphase (left images for all experimental conditions) and mitotic cells (right images). (B) SW13/cl.2 cells transfected with RFP//vimentin wt or (1-411) were treated in the absence or presence of 5 µM nocodazole for 30 min in serum-free medium. Vimentin (green) and tubulin (red) were visualized by immunofluorescence. (C) SW13/cl.2 cells were transfected as above and fixed. F-actin was stained with Phalloidin (red). Fluorescence intensity profiles of vimentin and actin along the dotted lines are shown in the right panels. (D) SW13/cl.2 cells transfected with RFP//vimentin wt or (1-411) were cultured in the absence or presence of 10 µg/ml cytochalasin B for 30 min in serum-free medium. Vimentin and f-actin were detected as above and their fluorescence intensity profiles along the dotted lines are shown in the right panels. (E) Cells transfected with RFP//vimentin wt were treated in serum-free medium with 2.5 µM latrunculin A for 30 min, 2 µg/ml C3 toxin for 3 h, 50 nM jasplakinolide for 30 min, or 20 µM blebbistatin for 1 h, as indicated, and processed by immunofluorescence. Images of single sections taken at mid-height of resting and dividing cells are shown.

Vimentin wt shows little coincidence with filamentous actin (f-actin) in interphase (Fig. 3C). In mitosis, f-actin accumulates at the cell periphery forming the actomyosin cortex, a stiff structure that allows spindle formation and orientation and maintains the cells spherical shape 34,35. Interestingly, peripherally distributed vimentin appeared to line the internal surface of the actomyosin cortex, partially overlapping with actin (Fig. 3C, fluorescence intensity profiles). Conversely, vimentin(1-411) did not follow the actin pattern under any condition (Fig. 3C). Importantly, vimentin cortical association was not a mere consequence of cell rounding, since newly-plated round cells showed vimentin perinuclear distribution, clearly unrelated to actin (Supplementary Fig. 2).

Although vimentin and actin show a reciprocal regulation at several cellular structures in resting cells ^36,37^, their interplay in mitosis has not been explored. Disruption of actin polymerization with cytochalasin B (Fig. 3C) B elicited a patchy f-actin distribution without severely affecting vimentin wt in resting cells. In mitotic cells, vimentin appeared in bundles entangled with dividing chromosomes, resembling vimentin(1-411), which was not further altered by cytochalasin B. Latrunculin A markedly decreased f-actin, leading to scattered aggregates (Fig. 3E). Loss of f-actin in mitosis correlated with vimentin bundling and intertwining with chromosomes. Interestingly, treatment with C3 toxin to inhibit Rho proteins, which are important for actomyosin cortex assembly^38,39^, resulted in a less homogeneous mitotic cortex and compromised vimentin peripheral distribution. Lastly, jaspalkinolide ^40^ elicited actin aggregates co-localizing with vimentin filaments in resting cells, and partially disrupted cortical association of vimentin in mitosis (Fig. 3E). In non-dividing cells, the myosin II ATPase inhibitor blebbistatin ^41,42^ induced a spikier actin pattern and intense shrinking of cell margins, where vimentin was condensed. In mitotic cells, the actin cortex was more irregular and vimentin was partially dislodged from the cell periphery (Fig. 3E). These results support an important role of f-actin and mitotic cortex integrity in the tail-dependent mitotic redistribution of vimentin.

### 4. Cortical localization of vimentin occurs in several cell types and is altered under pathophysiological conditions

Next, we explored the association of vimentin with the mitotic cortex in several cell types. In MCF7 cells, transfected vimentin showed signs of disassembly in mitosis and limited cortical association (Fig. 4A). In contrast, in astrocytoma and primary endothelial cells filamentous structures of endogenous vimentin adopted a peripheral distribution, close to the actin cortex (Fig. 4A). Primary human fibroblasts showed a rim of cortical vimentin together with loose vimentin bundles that did not interfere with chromosomes (Fig. 4A and Supplementary Fig. 3). Therefore, cortical distribution of vimentin in mitosis is cell-type dependent.

**Fig. 4.**
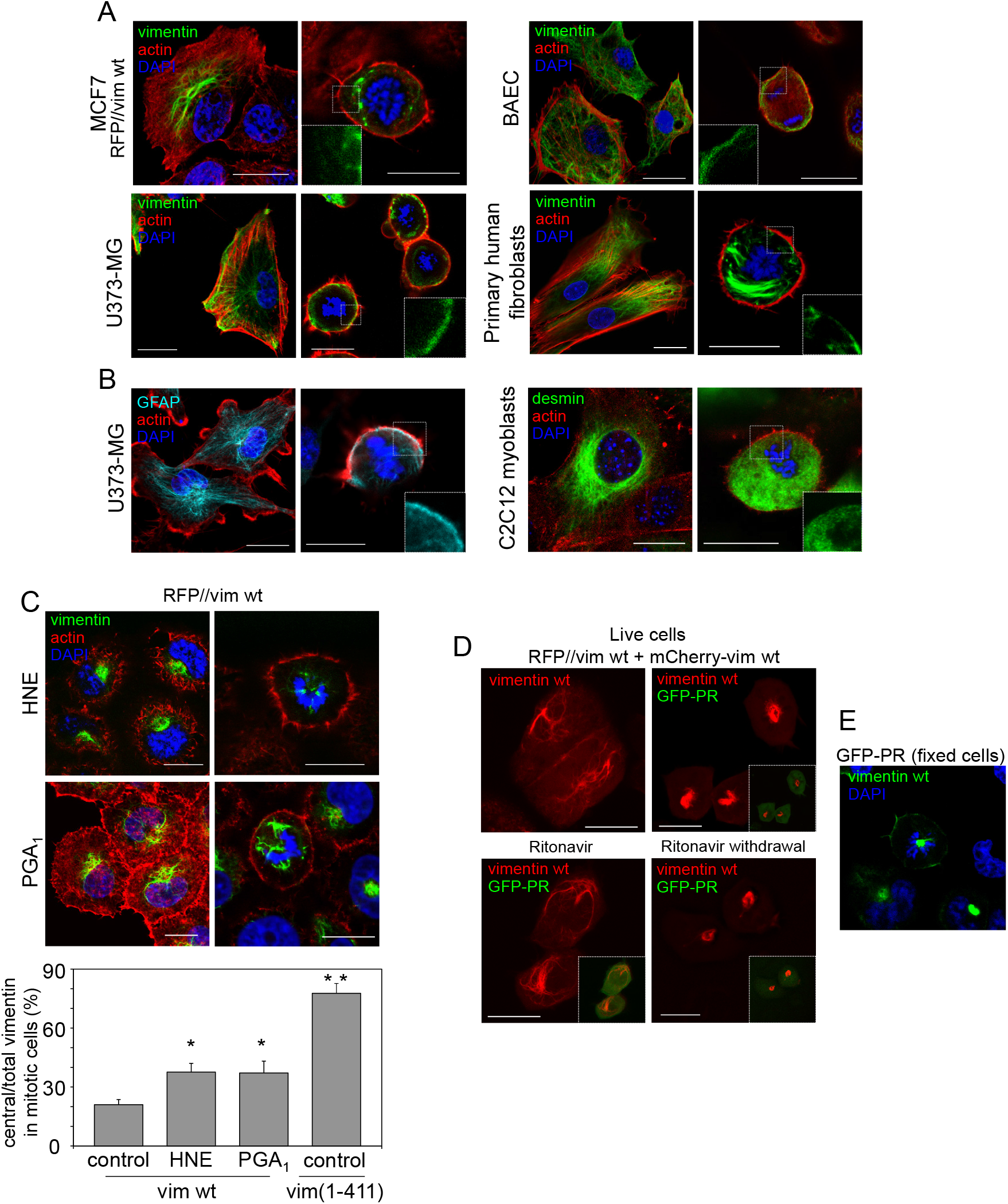
Association of type-III intermediate filaments with the mitotic cortex in several cell types and pathophysiological conditions. (A) The distribution of f-actin and vimentin in interphase and mitotic cells was assessed in several cell types expressing endogenous (astrocytoma U-373 MG cells, bovine aortic endothelial cells (BAEC), human dermal fibroblasts) or transfected (breast carcinoma MCF7 cells) vimentin. Inserts depict enlarged areas of the cell periphery showing vimentin distribution. Nuclei were counterstained with DAPI. Single sections at mid-cell height are shown in all panels. Single channel images, as well as fluorescence intensity profiles for actin and vimentin, are shown in Supplementary Fig. 3. (B) The distribution of GFAP and desmin was assessed by immunofluorescence in mitotic human astrocytoma U-373 MG cells and murine C2C12 myoblasts, respectively. (C) SW13/cl.2 cells expressing RFP//vimentin wt were treated with electrophilic lipids, 10 µM 4-hydroxynonenal (HNE) for 4 h or 20 µM prostaglandin A_1_ (PGA_1_) for 20 h, and processed as above. The histogram shows average values ± SEM from 15 determinations of the central versus total vimentin cellular fluorescence ratio, as an index of the impairment of peripheral localization. The values corresponding to vimentin(1-411) are included here for comparison. \**P*<0.05; \*\**P*<0.001 by Student’s *t*-test. (D) SW13/cl.2 cells were transfected with a combination of RFP//vimentin wt (80%) plus a tracer amount of mCherry-vimentin wt (20%), to monitor vimentin filaments (red) in live cells. In addition, cells were transfected with GFP-PR, coding the HIV-type I protease, and the corresponding green fluorescence is depicted in insets. In upper panels, cells were imaged 24 h after transfection. In the lower panels, the HIV inhibitor ritonavir was added immediately after transfection and cells were imaged 24 h later (left panel). Note the more intense GFP fluorescence indicative of the inhibition of GFP-PR autolysis. Subsequently, ritonavir was removed and cells were imaged 5 h later (right panel). (E) At the end of the experiment, cells were fixed and stained with DAPI to identify mitotic cells. Vimentin is artificially colored in green.

Importantly, filaments of glial fibrillary acidic protein (GFAP), another type III intermediate filament protein, also adopted a peripheral distribution in mitotic astrocytoma cells, whereas desmin showed a predominantly diffuse cytoplasmic staining in mitotic C2C12 myoblasts (Fig. 4B).

We then assessed the distribution of vimentin filaments in the presence of pathophysiological agents known to cause vimentin “collapse”. The inflammatory lipid mediators 4-hydroxynonenal and prostaglandin A_1_ induced vimentin juxtanuclear condensation in interphase cells, as previously observed ^12^. Moreover, they significantly dislodged vimentin from the mitotic actomyosin cortex (Fig. 4C), according to the central/total vimentin fluorescence ratio (Fig. 4C). Additionally, expression of a HIV-protease construct (GFP-PR) was associated with collapse of vimentin filaments into curly juxtanuclear bundles (Fig. 4D), reminiscent of the vimentin(1-411) distribution. Ritonavir, a reversible HIV-protease inhibitor, blocked vimentin collapse, whereas its withdrawal allowed fast vimentin condensation, indicating a role for protease activity (Fig. 4D). HIV-protease-induced vimentin accumulations remained in mitosis and concentrated close to the dividing chromosomes (Fig. 4E). These results show that various pathophysiological agents cause anomalous vimentin distribution in mitosis, hampering cortical localization.

### 5. Vimentin is intimately intertwined with actin at the mitotic cortex

Analysis of the interaction of vimentin with the actomyosin cortex by superresolution microscopy (STED) showed vimentin filaments next to the cortex, intermingling with actin at some points (Fig. 5A, upper two rows) or running between two actin layers (Fig. 5A, lower two rows). Single section analysis of actin-vimentin co-localization showed a closer connection at certain locations along the mitotic cortex, suggesting the existence of docking or penetration sites of vimentin in this structure. Three-dimensional reconstructions revealed a robust basket-shaped framework of vimentin filaments of diverse orientations (Fig. 5B). Actin formed a hollow sphere constituted by elongated patches or bundles mostly oriented perpendicularly to the support surface, as illustrated in the 3D-reconstruction of the cell bottom half (Fig. 5C and Supplementary video 5). Interestingly, this reconstruction evidences points of vimentin filament protrusion through the actin cortex (Fig. 5C).

**Fig. 5.**
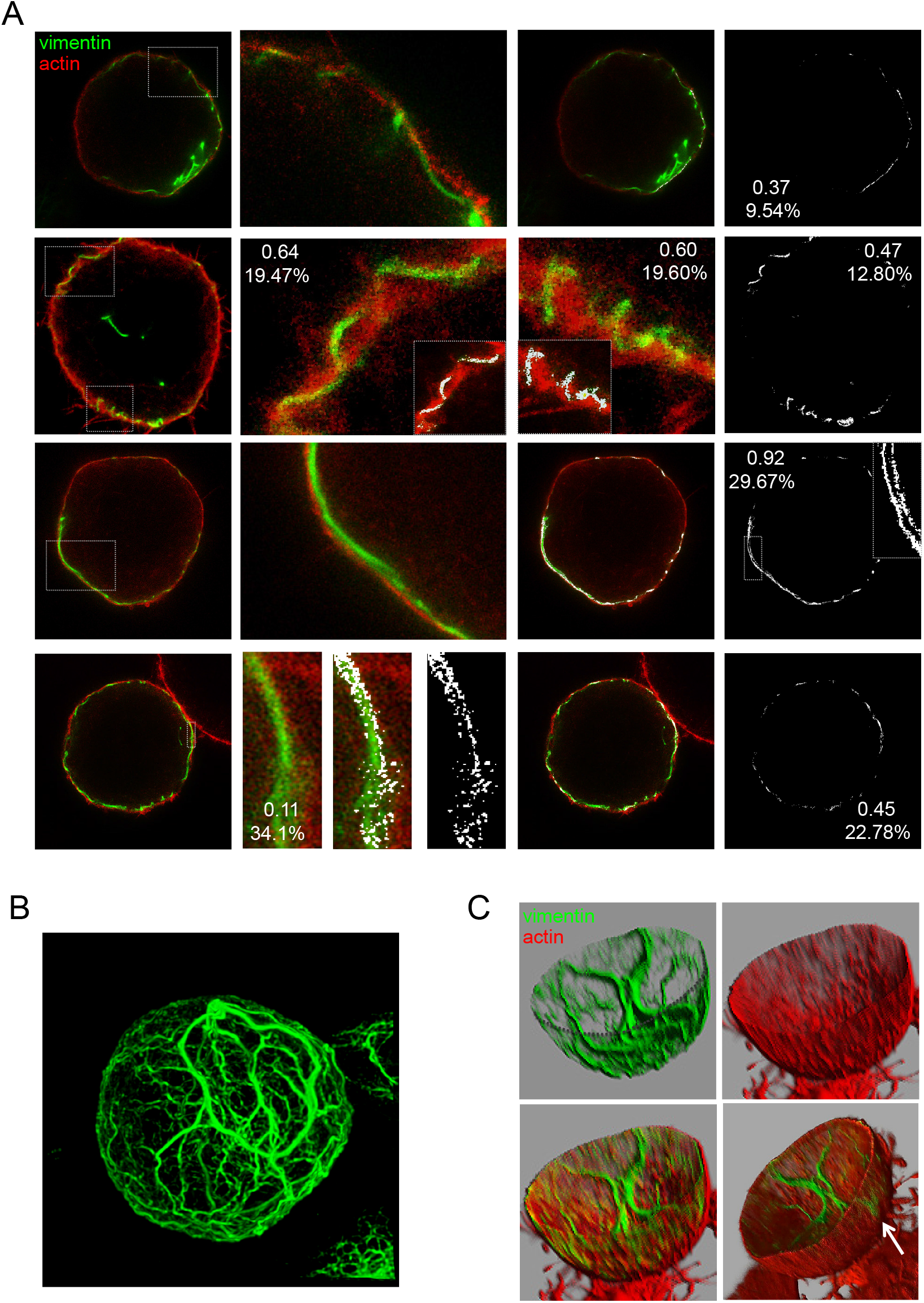
Analysis of the relative positions of vimentin and actin in mitosis by STED superresolution microscopy. SW13/cl.2 cells stably transfected with RFP//vimentin wt were treated with 0.4 µM nocodazole overnight, to increase the proportion of cells in mitosis, as described in the experimental section. Vimentin was detected by immunofluorescence with Alexa488-conjugated V9 antibody and actin was stained with TRITC-Phalloidin. (A) STED images of several cells are shown. Co-localization analysis was performed with Leica software. Numbers in insets represent the Pearson’s coefficient and the percentage of co-localization, respectively, for the whole cell or for the regions enlarged, as indicated. (B) 3D-reconstruction of vimentin organization, after deconvolution of the green channel using Imaris software, for one representative cell. (C) 3D-reconstruction using the basal half of the sections from the same cell in order to show the “inside” and the “outside” of the sphere. Single channels (upper panels) and merged images (lower panels) are shown. The semi-sphere edge is marked in the green channel (dotted line). The bottom-right image is a snapshot of Supplementary video 5. A point where vimentin protrudes through the actin cortex is indicated by an arrow.

### 6. Vimentin-actin interplay in mitotic cells

Deeper insight into the vimentin-actin interaction at the mitotic cortex was obtained by the analyses schematized in Figure 6A. First, we obtained 3D-reconstructions of the cortex of vimentin-positive and -negative cells, which confirmed that some cells displayed ample segments of vimentin at the external surface of the cortex, interwoven with actin structures (Fig. 6B and Supplementary videos 6 and 7). Next, 2D-map projections from the image stacks were prepared for global visualization and quantitation of the cortex ^43^ (Fig. 6B). Fluorescence intensity profiles of these projections illustrated the alternate distribution of actin and vimentin signals at some points. 2D-maps from vimentin-negative cells presented higher standard deviation of f-actin pixel brightness, indicating wider variations in f-actin distribution (Fig. 6B). Additionally, we analyzed orthogonal projections of vimentin-positive and -negative mitotic cells (Fig. 6C). Notably, vimentin filaments could be detected both at the top and bottom of vimentin-positive cells (Fig. 6C, arrowheads), with a particular enrichment of robust lattices at the basal layer, next to the substrate. These structures were obviously absent from non-transfected cells, but also from cells expressing vimentin(1-411), which was frequently retained close to the inner actin “ring-like” structure (Fig. 6C, inset), clearly detectable in some cells, which has been involved in spindle positioning ^44^. Additionally, the basal vimentin lattice was associated with a decreased f-actin signal and lower standard deviation of pixel brightness at this location, suggestive of less polymerized actin structures (Fig. 6C, graph). Altogether, these results indicate that vimentin may be an important player at the mitotic cortex, exerting a measurable impact on its characteristics that could influence cell division dynamics.

**Fig. 6.**
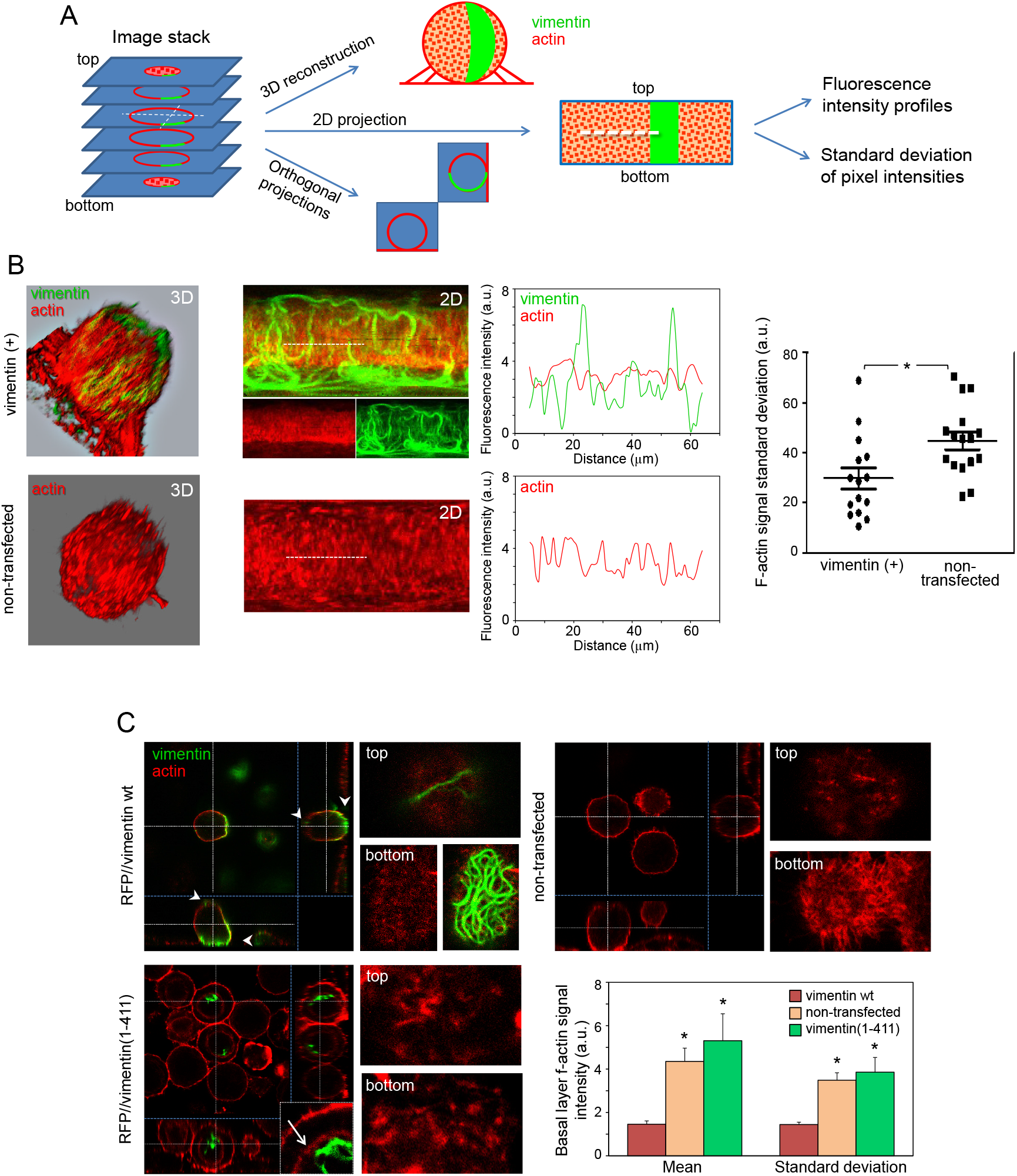
Vimentin expression affects the properties of the mitotic cortex. (A) Scheme of the different approaches employed to analyze mitotic cell properties. XY image stacks were used to obtain 3D-reconstructions of mitotic cells, 2D “cartographic”, or orthogonal projections. (B) From left to right, 3D-reconstruction of SW13/cl.2 cells, stably expressing vimentin wt (top) or non-transfected (bottom); 2D-maps from the same cells (for the vimentin-expressing cell, the merged and single channels are shown); fluorescence intensity profiles along the white dotted lines drawn on the 2D-maps; scattered plot of standard deviation values of the actin signal of 2D-maps from 16 cells per experimental condition; \**P*< 0.05 by unpaired *t*-test. (C) SW13/cl.2 cells were transfected with RFP//vimentin wt (stable transfection), RFP//vimentin(1-411) (transient transfection) or non-transfected. Orthogonal projections illustrate the positions of the vimentin constructs. Arrowheads mark the appearance of vimentin wt at the top and at the bottom of the cell. The inset shows an enlarged image illustrating the localization of vimentin(1-411) with respect to the cytoplasmic actin ring (marked by an arrow). Right panels show the top and bottom sections for every construct. The histogram depicts the mean and standard deviation of f-actin signal intensity at the bottom section. Values are average ± SEM from at least 10 determinations. \**P*< 0.05 vs vimentin wt.

### 7. Impact of serial C-terminal deletions on vimentin organization and mitotic peripheral distribution

Structural determinants allowing vimentin to reach the cell periphery in mitosis were explored by analyzing the distribution of mutants bearing several C-terminal deletions (Fig. 7A). Vimentin(1-423), mimicking the reported HIV-protease cleavage product ^23,45,46^, formed curly bundles in the nuclear vicinity (Fig. 7B), similar to vimentin(1-411). Vimentin(1-423) coiled bundles mainly remained near the condensed chromosomes in mitosis, either interfering with the mitotic spindle or located at one of the poles (Fig. 7C). Often, multi-nucleated cells containing coiled vimentin(1-423), and in some cases DNA, in the space between nuclei were found (Supplementary Fig. 1), suggesting cytokinetic defects. Time-lapse monitoring of cells expressing vimentin(1-423) confirmed marked mitotic alterations, including vimentin asymmetric partitioning, delayed mitosis and cell death (Fig. 7D and Supplementary videos 8 and 9). Moreover, vimentin(1-423), was often retained at the cytoplasmic f-actin ring and did not reach cortical actin (Fig. 7E).

**Fig. 7.**
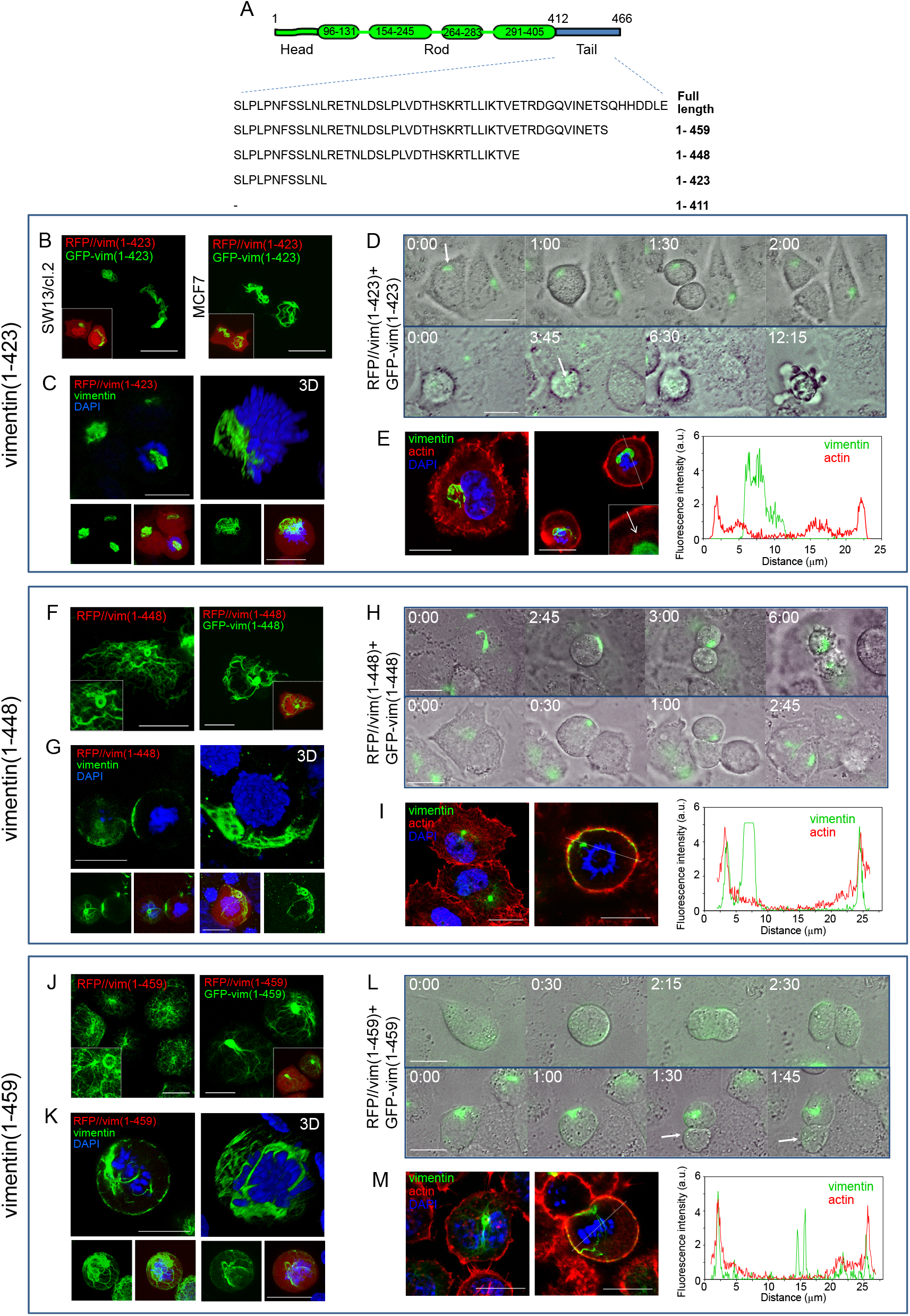
Organization and mitotic distribution of several C-terminal truncated vimentin mutants. (A) Scheme of the various vimentin truncated mutants generated. (B) Overall projections of live SW13/cl.2 or MCF7 cells transfected with RFP//vimentin(1-423) plus GFP-vimentin(1-423). Insets depict the overlay with the RFP background fluorescence delimiting the cell contour. (C) SW13/cl.2 cells were transfected with RFP//vimentin(1-423) and the distribution of truncated vimentin in mitosis was assessed by immunofluorescence. Nuclei were counterstained with DAPI. A representative single section at mid-cell height (left) and a 3D-projection (right) are shown. Lower panels depict overall projections of vimentin alone (green channel) or the overlay of vimentin, DAPI and the red background fluorescence. (D) Live cells transfected with RFP//vimentin(1-423) plus GFP-vimentin(1-423) were monitored by time-lapse microscopy. Representative cases of vimentin asymmetric partition (upper panels) and mitotic catastrophe (lower panels) are shown. (E) Cells were transfected as in (C) and the distribution of vimentin and f-actin was monitored in interphase (left) and mitosis (right). Images are single sections at mid-cell height. The arrow marks the position of the cytoplasmic actin ring. The right panel shows fluorescence intensity profiles of vimentin and actin signals along the dotted line drawn on a mitotic cell. (F and J) Cells were transfected with RFP//vimentin(1-448) in (F), or RFP//vimentin(1-459) in (J), alone or in combination with the corresponding GFP fusion construct, as indicated, and vimentin distribution was monitored by immunofluorescence (left panels) or live cell direct visualization (right panels). (G and K) Cells transfected with RFP//vimentin(1-448) in (G), or RFP//vimentin(1-459) in (K) were analyzed as in (C). (H) Representative sequence from time-lapse monitoring of live cells transfected with RFP//vimentin(1-448) showing an incomplete division ending in cell death (upper panel) and an asymmetric division (lower panel). (L) Sequence from live-cell monitoring after transfection with RFP//vimentin(1-459) showing a normal division (upper panel) and an asymmetric division (lower panel). (I and M) The distribution of vimentin and f-actin in cells transfected with RFP//vimentin(1-448) (I) or RFP//vimentin(1-459) (M) was monitored in resting and mitotic cells, as described for panel (E).

Vimentin(1-448) (Fig. 7A), yielded a heterogeneous pattern with both extended filaments and robust bundles or accumulations (Fig. 7F). These persisted in mitotic cells, sometimes appearing at basal planes or at the cell periphery (Fig. 7G). Mitotic cells suffered two main fates: approximately 40% exhibited delayed separation ending in cell death, indicative of cytokinetic failure, whereas 60% completed mitosis through vimentin asymmetric partition (Fig. 7H and supplementary videos 10 and 11). Nevertheless, some peripheral filamentous vimentin could be detected, lying adjacent to the actomyosin cortex (Fig 7I).

Finally, a construct with a shorter C-terminal deletion, vimentin(1-459), formed filaments similar in morphology and extension to those of vimentin wt, although ∼18% of the cells also showed small bundles or curls (Fig. 7J). In mitosis, vimentin(1-459) adopted a mainly peripheral distribution, although some cells presented filaments intertwined with chromosomes (Fig. 7K). In time-lapse monitoring (Fig. 7L and supplementary videos 12 and 13), cells lacking vimentin bundles underwent basically normal mitosis with even vimentin distribution between daughter cells. Conversely, cells harboring vimentin bundles showed a partial asymmetric distribution, with one daughter cell receiving most of vimentin(1-459) (Fig. 7L). Vimentin(1-459) coincided with some segments of the actomyosin cortex, whereas some filaments persisted in the central area (Fig. 7M). Thus, deletion of the last seven amino acids induces a mild perturbation of vimentin distribution in mitosis.

Taken together, these results reveal that step-wise deletion of the tail gradually impairs normal vimentin assembly and redistribution in mitosis, with abolishment of mitotic peripheral localization being observed upon vimentin truncation at L423 or I411.

### 8. Vimentin cortical association in mitosis does not require network formation or full filament assembly

To discard that lack of cortical association could be due to intense bundling, we used GFP-vimentin fusion constructs, which do not form full filaments in SW13/cl.2 cells ^12^. First, the organization of GFP-vimentin wt and all the truncated variants was studied (Fig. 8A). GFP-vimentin wt formed a uniform lattice of squiggles or short filaments ^12^ (Fig 8A). Conversely, GFP-vimentin(1-411) could not reach the squiggle stage and formed only bright dots. GFP-vimentin(1-423) exhibited a mixed pattern consisting of dots, small swirls and occasional short filaments. Longer constructs, namely, GFP-vimentin(1-448) and GFP-vimentin(1-459) often presented short filaments or squiggles (Fig. 8A). Thus, GFP-fusion constructs displayed a gradual sequence-dependent impairment of particle elongation in vimentin-deficient cells, stressing that even small truncations of the tail domain have a detectable impact (quantitated in Fig. 8, graph).

**Fig. 8.**
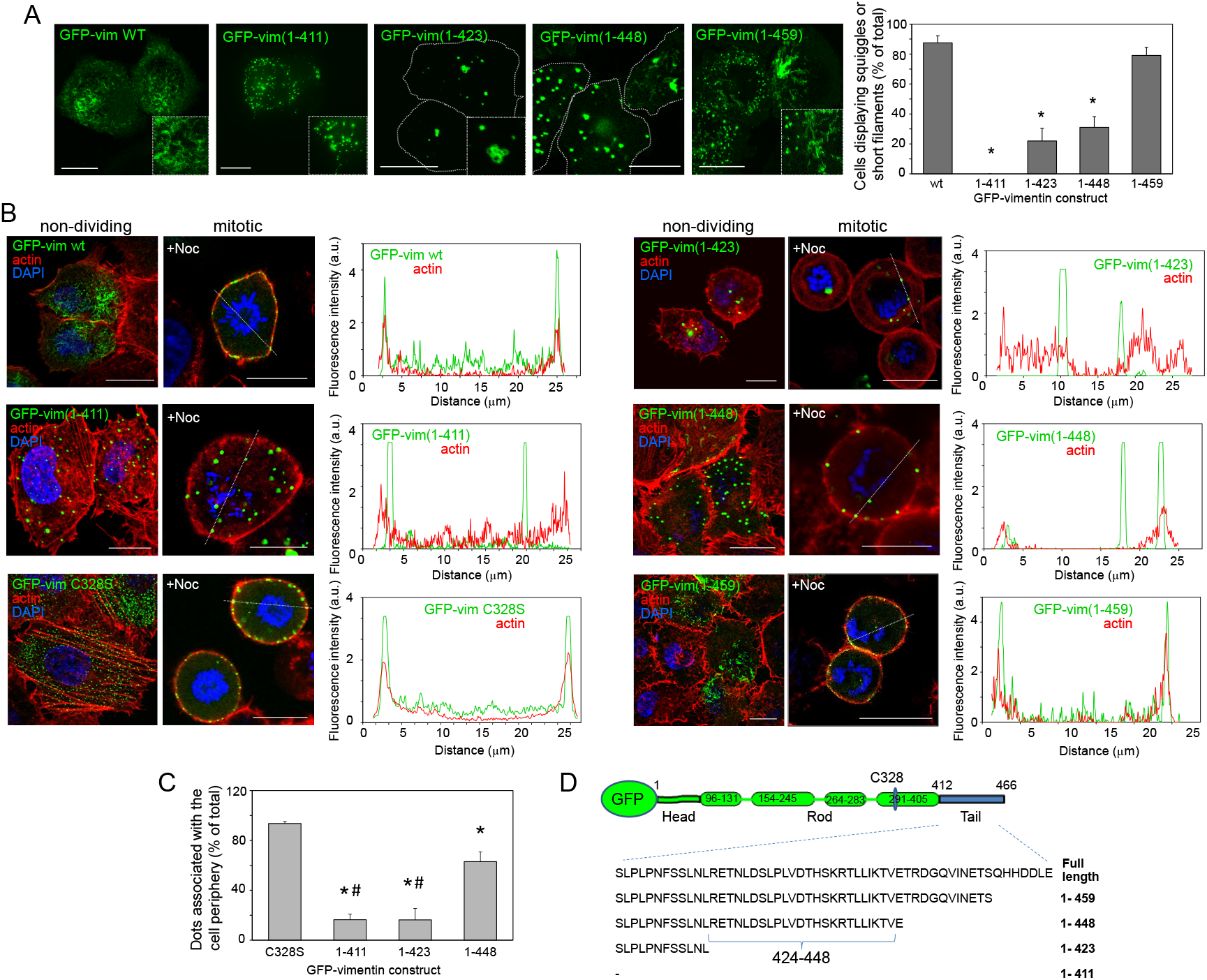
Assembly and distribution of GFP fusion constructs of truncated forms of vimentin. (A) Live confocal microscopy assessment of the morphology of vimentin assemblies 48 h after transfection of SW13/cl.2 cells with the constructs shown in panel (D). Overall projections are shown. Insets show enlarged areas of interest. The histogram (right) depicts the percentage of cells with squiggles or short filaments for every construct. Results are average values ± SEM of at least 15 fields from several experiments totaling at least 100 cells per experimental condition.\**P*<0.05 by Student’s *t*-test. (B) Cells were transfected with the indicated constructs, fixed, and the distribution of vimentin and f-actin analyzed as above. Both interphase (left images) and mitotic cells (right images) are shown. Where indicated, cells were treated overnight with 0.4 µM nocodazole (+Noc) in complete medium to increase the proportion of mitotic cells. Nuclei were counterstained with DAPI. Single sections taken at mid-cell height are shown in all cases. The fluorescence intensity profiles for vimentin and f-actin along the dotted lines are shown on the right for every condition. (C) The histogram shows the proportion of dots associated with the cell periphery for every construct. In this case, GFP-vimentin C328S, which contains the tail domain but assembles in dots, is used as a control. GFP-vimentin(1-459) is excluded from this graph due to its organization mainly in squiggles or short filaments. Results are average values of at least 10 fields from at least three experiments per condition. \**P*<0.01 vs GFP-vimentin C328S; #*P*<0.001 vs GFP-vimentin(1-448). (D) Scheme illustrating the sequence of the various GFP-vimentin truncated constructs. The approximate position of C328 is indicated.

Most constructs showed little overlap with actin structures in resting cells, with GFP-vimentin wt, C328S and GFP-vimentin(1-459) displaying more points of contact (Fig. 8B). In cells arrested in mitosis by mild nocodazole treatment, GFP-vimentin constructs frequently showed a diffuse background, suggestive of a higher extent of disassembly than untagged vimentin. Nevertheless, GFP-vimentin wt structures were clearly detected at the mitotic cell cortex co-localizing with actin (Fig. 8B). In sharp contrast, GFP-vimentin(1-411) dots appeared scattered throughout the cell. This lack of cortical association cannot be solely attributed to defective elongation, since dots formed by full-length GFP-vimentin C328S, which is also elongation-incompetent ^12^, relocated to the periphery of mitotic cells (Fig 8B). GFP-vimentin(1-423) accumulations also failed to associate with the actin cortex, frequently appearing near the cytoplasmic f-actin ring, whereas > 60% of GFP-vimentin(1-448) dots redistributed to the cell periphery and cortical localization of GFP-vimentin(1-459) mixed structures was preserved (Fig. 8C, graph). Thus, serial tail truncations exert a graded impact on mitotic cortical association, denoting the importance of the segment 424-448 for this redistribution (Fig. 8D). Moreover, full filament elongation is not necessary for cortical association since particles formed by constructs retaining all or most of the tail domain, effectively relocate to the cell periphery. Nevertheless, a certain degree of assembly seems necessary since neither GFP-vim(412-466) (completely diffuse) nor the assembly-incompetent vimentin∆3-74 mutant were able to redistribute to the mitotic cortex (Supplementary Fig. 4).

Thus, the tail domain is essential, but not sufficient, for vimentin cortical association in mitosis, and other structural or conformational factors appear necessary.

## Discussion

Vimentin plays critical functions in cell mechanics ^47^. Nevertheless, its role in mitosis is not fully understood. Here we unveil the robust scaffold formed by vimentin filaments in mitosis in several cell types. This framework intimately interacts with the actomyosin cortex, intertwining with actin, and affecting its properties. Several functions can be envisaged for this arrangement: to yield space for mitotic spindle organization, and, potentially, to modulate the robustness or stiffness of the mitotic cortex. Therefore, these results warrant the pertinence to study vimentin, and other intermediate filaments, as players in mitotic cortex dynamics.

Vimentin organization in mitosis is cell-type dependent and responds to two main patterns: formation of a filament “cage” surrounding the mitotic spindle, or disassembly induced by phosphorylation of N-terminus residues in combination with protein-protein interactions, reportedly, copolymerization with nestin ^48,49,50^. We observed vimentin rearrangements potentially related to nestin levels, with SW13/cl.2 cells (nestin-negative) retaining filaments, MCF7 cells (nestin-positive) ^51^, showing vimentin disassembly, and mixed patterns in the other cell types which express variable nestin:vimentin proportions ^52^. It can be hypothesized that, if not disassembled, vimentin filaments should undergo mitotic cortical translocation or anchorage to facilitate mitosis progression. This is substantiated by the striking behavior of vimentin(1-411), which does not reach the actomyosin cortex and interferes with the mitotic apparatus causing aberrant mitosis. This raises potential cytotoxic implications of vimentin cortical dislodgement in pathophysiological settings, as observed upon lipoxidation or C-terminal cleavage, although damage or cleavage of macromolecules different than vimentin could contribute to these effects.

The intrinsically disordered C-terminal vimentin domain has been proposed to undergo conformational rearrangements during filament assembly and to participate in protein-protein interactions, including actin ^53,54^. Our results indicate that in mitosis, filamentous vimentin interacts with the actomyosin cortex showing points of co-localization with actin. However, this interaction could take place through other proteins, including the scaffold protein plectin, chaperones or actin-associated proteinst^55^. Additionally, vimentin protrudes through the actomyosin cortex at some points, for which sites of attachment at the plasma membrane involving protein receptors or lipid domains, cannot be excluded.

A complex interplay between actin and vimentin at several organization levels exists in resting cells ^36^. Actin limits transport of vimentin ULF along microtubules ^56^, and actomyosin arcs interact with vimentin filaments through plectin and drive their retrograde movement, thus promoting vimentin perinuclear localization ^37^. In turn, vimentin restricts retrograde movement of the arcs and restrains actin polymerization and stress fiber assembly ^57^. Nevertheless, the actin-vimentin interaction in mitosis has not been addressed to our knowledge. The mitotic cortex provides tension, which together with osmotic pressure controls cell rounding. Actin organization is a key factor in contractile tension generation ^58^. Importantly, our results clearly show that integrity of the actomyosin cortex is necessary for vimentin cortical association, but also suggest an impact of vimentin on cortical actin organization. The lower dispersion of f-actin signal in global cortex 2D-projections and the lower intensity at basal planes in vimentin-positive cells suggests a negative feedback on the formation of highly polymerized actin structures, which in mitotic SW13/cl.2 cells appear mainly as elongated actin bundles perpendicular to the substrate. Thus, our studies open the way for dissecting the consequences of vimentin-actin interplay on actomyosin contractility or stiffness in mitosis ^59^.

Vimentin tail integrity is determinant for cortical association. On one hand, both untagged and GFP-fusion constructs show a graded impairment of cortical association upon serial tail truncations, the strongest impact occurring after deletion of the 43 distal residues. Therefore, the sequence comprised between residues 424-448, appears to contain important determinants for mitotic redistribution. This segment approximately coincides with a putative loop proposed to protrude from filaments and participate in protein-protein interactions ^60,61^. Although this contention still needs conclusive experimental evidence, our results raise the interest of investigating posttranslational modifications affecting the vimentin tail, which could modulate cortical association. On the other hand, although vimentin polymerization into full filaments is not necessary for cortical localization, a certain level of organization appears necessary since soluble vimentin forms do not undergo cortical association, for which integrity of the N-terminus could also play a role.

Thus, our results, summarized in Supplementary Figure 8, show that vimentin filaments redistribute to the cell periphery in mitosis in a tail domain-dependent manner. This reorganization implies a close interplay with the actomyosin cortex, which sets forth novel functions of intermediate filament dynamics during the cell cycle and opens the way for the search of strategies modulating these interactions.

## Materials and methods

Reagents. Restriction enzymes and buffers were from Promega. Anti-vimentin antibodies were: mouse monoclonal V9 clone (sc-6260) and its Alexa-488 conjugate from Santa Cruz Biotechnology, and mouse anti-vimentin monoclonal antibody (V5255) from Sigma. Anti-actin (A2066) was from Sigma and anti-α-tubulin (ab52866) and anti-desmin (ab15200-1) from Abcam. Anti-GFAP (Z0334) was from Dako. C3 transferase toxin was from Cytoskeleton. Latrunculin A and jasplakinolide were from Santa Cruz Biotechnology. 4-hydroxynonenal (HNE) and prostaglandin A_1_ (PGA_1_) were from Cayman Chemical. 4,6-diamidino-2-phenylindole (DAPI), blebbistatin and ritonavir were from Sigma.

Cell culture and treatments. SW13/cl.2 human adrenocarcinoma vimentin-deficient cells were the generous gift of Dr. A. Sarriá (University of Zaragoza, Spain)^62^. MCF7 human breast carcinoma cells, U-373 MG human glioblastoma astrocytoma cells and C2C12 murine myoblasts were from ATCC. They were cultured in DMEM with 10% (v/v) fetal bovine serum (FBS) and antibiotics (100U/ml penicillin and 100μg/ml streptomycin). Bovine aortic endothelial cells (BAEC) were from Clonetics and were cultured in RPMI1640 supplemented with 10% (v/v) newborn calf serum (Gibco) and antibiotics. Primary human dermal fibroblasts from an adult donor (ref. AG10803) were obtained from the NIA Aging Cell Repository at the Coriell Institute for Medical Research (Camden, NJ). Unless otherwise stated, treatments were carried out in serum-free medium. For acute microtubule disruption, cells were treated with 5 µM nocodazole for 30 min. For mitotic arrest, cells were cultured in the presence of 0.4 µM nocodazole for 20 h in complete medium. This treatment was employed, when indicated, to increase the proportion of mitotic cells in conditions under which neither actin, nor vimentin organization were altered with respect to untreated cells (Supplementary Fig. 5). Disruption of f-actin was achieved by treatment with 10 µM cytochalasin B or 2.5 µM latrunculin A for 30 min, or 2 µg/ml C3 toxin for 3.5 h. Jasplakinolide was employed at 50 nM for 30 min. Blebbistatin was used at 20 µM for 1 h. For treatment with electrophilic lipids, cells were incubated in the presence of 10 µM HNE for 4 h or 20 µM PGA_1_ for 20 h. Inhibition of transfected HIV protease was achieved by incubating the cells in the presence of 10 µM ritonavir in serum-containing medium, immediately after transfection. For removal of the inhibitor cells were washed three times with fresh medium with serum, without antibiotics.

Plasmids and transfections. The bicistronic plasmid RFP//vimentin wt, coding for the red fluorescent protein DsRed Express2, abbreviated as RFP, and wt human vimentin as separate products, and the GFP fusion constructs, GFP-vimentin wt and GFP-vimentin C328S have been previously describedt^12,63^. mCherry-vimentin was from Genecopoeia. The various tail truncated mutants, vimentin(1-411), (1-423), (1-448) and (1-459) were generated introducing stop codons at positions 412, 424, 449 and 460, respectively, by site directed mutagenesis of the parent vectors using the Quikchange XL mutagenesis kit from Stratagene and the primers specified in Supplementary Table 1, following the instructions of the manufacturer. Truncated vimentin constructs showed the expected mobility in SDS-PAGE gels as well as immunoreactivity (Supplementary Fig. 6A). Thus, all constructs used were recognized by an antibody raised against full-length vimentin, which recognizes the N-terminus of the protein (anti-vim N-term), whereas an antibody against the beginning of the tail (clone V9) recognized all constructs except tailless vimentin(1-411). The GFP-vimentin(412-466) construct, encoding the vimentin tail domain fused with GFP, was constructed in two steps; first, an additional EcoRI site was introduced in the GFP-vimentin wt plasmid (containing the vimentin sequence cloned between the EcoRI and BamHI sites) at a position equivalent to 1692 of the vimentin mRNA sequence (accession number NM_03380.4); then, the plasmid was digested with EcoRI and re-ligated, using the Ligafast system from Promega, thus eliminating the sequence corresponding to vimentin residues 1 to 411. The assembly-incompetent construct RFP//vimentin∆3-74 was generated by introducing a SmaI site at a position equivalent to 471 of the vimentin mRNA sequence in the RFP//vimentin wt plasmid through site-directed mutagenesis. The resulting mutant plasmid was digested with SmaI and re-ligated resulting in the removal of nucleotides 472 through 687, thus eliminating residues 3 to 74. The CFP-lamin A plasmid was the gift of Dr. Vicente Andrést ^64^. The vector encoding GFP-tagged HIV type I protease (pcDNA3/GFP-PR) described in ^65^ was a gift from Nico Dantuma (Addgene plasmid #20253). Cells were transfected using Lipofectamine 2000 (Thermo Scientific), as previously described ^12,66^. Typically, 1 µg of DNA and 3 µl of Lipofectamine 2000 were used per p35 dish. For overexpression of RFP//vimentin wt or (1-411) in U-373 MG astrocytoma cells, 2 µg of DNA plus 4.5 µl of Lipofectamine 2000 were used. For expression of different proportions of vimentin wt and tailless (1-411), the following plasmid amounts were used: 10:0, 0.8 µg RFP//vim wt + 0.2 µg GFP-vim wt; 8:2, 0.8 µg RFP//vim wt + 0.2 µg GFP-vim(1-411); 4:6, 0.4 µg RFP//vim wt + 0.4 µg RFP//vim(1-411) + 0.2 µg GFP-vim(1-411); 2:8, 0.2 µg RFP//vim wt + 0.6 µg RFP//vim(1-411) + 0.2 µg GFP-vim(1-411). Routinely, cells were visualized 48 h after transient transfection. When indicated, cells were cultured in the presence of 500 µg/ml G-418 for generation of stably transfected cells.

Fluorescence microscopy and image analysis. Cells transfected with the various constructs were visualized live by confocal microscopy on Leica SP2 or SP5 microscopes. Images were acquired every 0.5 µm and single sections or overall projections are shown, as indicated. All scale bars are 20 µm. For immunofluorescence, cells were fixed with 4% (w/v) paraformaldehyde for 25 min at r.t., permeabilized with 0.1% (v/v) Triton-X100 in PBS and blocked with 1% (w/v) BSA in PBS. Antibodies were used at 1:200 dilution in blocking solution. For experiments involving detection of vimentin(1-411), the monoclonal antibody recognizing the vimentin N-terminus was used for all conditions. For experiments involving selective detection of full-length vimentin or not requiring a comparison with vimentin(1-411), the V9 antibody was employed (Supplementary Fig. 6B). F-actin was stained with Phalloidin-Alexa568 or Phalloidin-Alexa488 (Molecular Probes), following the manufacturer instructions. Nuclei were counterstained with DAPI (3 µg/ml). Direct visualization on glass-bottom culture dishes was found optimal for imaging mitotic cells, since mounting on glass slides compromised their spherical shape. For superresolution microscopy through stimulated emission depletion (STED), vimentin was detected with Alexa488-conjugated anti-vimentin V9 and f-actin was stained with Phalloidin-Tetramethylrhodamine B isothiocyanate from Sigma (0.25 µg/ml). Images were acquired with a confocal multispectral Leica TCS SP8 system equipped with a 3X STED module. Co-localization was analyzed with Leica software. Time-lapse microscopy was carried out in a multidimensional microscopy system Leica AF6000 LX in a humidified 5% CO_2_ atmosphere at 37ºC. Typically, green fluorescence and differential interference contrast (DIC) images were recorded. 3D-reconstructions were obtained with Image J (FIJI), Imaris or Leica software. Fluorescence intensity profiles and measurements of mean fluorescence intensity and standard deviation of pixel brightness values, to illustrate the dispersion of f-actin intensity, were obtained with ImageJ. Orthogonal projections were obtained with Leica software. 2D maps from image stacks were obtained by FIJI and the free Map3-2D software developed by Sendra el al., ^43^ (http://www.zmbh.uniheidelberg.de//Central_Services/Imaging_Facility/Map3-2D.html), which unfolds surface information onto a single structurally connected map, using a “sphere” adjustment. For quantitation of vimentin reorganization induced by electrophilic lipids, the proportion of vimentin fluorescence present in the central area of mitotic cells (central circle of a diameter of 60% the total cell diameter in a single section at mid-cell height), with respect to the total area was measured as an indication of the impairment of peripheral distribution.

SDS-PAGE and western blot. Cells transfected with the various constructs were lysed in 20 mM Tris-HCl pH 7.5, 0.1 mM EDTA, 0.1 mM EGTA, 0.1 mM β-mercaptoethanol, containing 0.5% (w/v) SDS, 0.1 mM sodium orthovanadate and protease inhibitors (2 µg/ml each of leupeptin, aprotinin and trypsin inhibitor, and 1.3 mM Pefablock), and processed essentially as described ^67^. Briefly, protein concentration in lysates was determined by the bicinchoninic acid assay. Aliquots of lysates containing 30 µg of total protein were denatured in Laemmli buffer for 5 min at 95ºC and separated in 10 or 15% SDS-polyacrylamide gels. Gels were transferred to Immobilon-P membranes (Millipore) using a Tris-glycine methanol three-buffer system, as recommended by the manufacturer, on a semi-dry transfer unit (Transblot) from Bio-Rad. Membranes were blocked with 2% (w/v) low-fat powdered milk in T-TBS (Tris-HCl pH 7.5, 500 mM NaCl, 0.05% (v/v) Tween-20). Subsequently, membranes were incubated with primary antibodies at 1:500 dilution and horseradish peroxidase-conjugated secondary antibodies (Dako) at 1:2000 dilution. Proteins of interest were detected with the ECL system from GE Healthcare.

Statistical analysis. All experiments were repeated at least three times with similar results. All results are presented as average values ± SEM. Statistical analysis was performed with GraphPad Prism. Statistical differences were evaluated by the unpaired Student’s *t*-test and were considered significant when *P*<0.05, which is denoted in graphs by an asterisk. The significance levels for every experiment are given in the figure legends.

## Acknowledgements

This work has been funded by the European Union’s Horizon 2020 research and innovation program under the Marie Sklowdowska-Curie grant agreement number 675132, “Masstrplan” (http://cordis.europa.eu/project/rcn/198275_en.html), and by grants from the Spanish Ministerio de Economía y Competitividad (MINECO/FEDER, http://www.mineco.gob.es/portal/site/mineco/idi) SAF2015-68590R and from Instituto de Salud Carlos III/FEDER, RETIC Aradyal RD16/0006/0021. AVP is supported by the FPI Program from MINECO, reference: BES-2016-076965. Feedback from the EU COST Action CA15214 “EuroCellNet” is gratefully acknowledged.

We are indebted to MT Seisdedos and Dr. G Elvira from CIB, and Dr. S Gutiérrez from CNB, CSIC for help with confocal and superresolution microscopy, respectively. We thank Dr. Francisco J. Sánchez-Gómez and Prof. F.J. Cañada for helpful comments and discussion. The valuable technical assistance of MJ Carrasco is gratefully appreciated.

